# An Effective Model of HL-60 Differentiation

**DOI:** 10.1101/029066

**Authors:** Ryan Tasseff, Holly A. Jensen, Johanna Congleton, Andrew Yen, Jeffrey D. Varner

## Abstract

We present an effective model All-Trans Retinoic Acid (ATRA)-induced differentiation of HL-60 cells. The model describes a key architectural feature of ATRA-induced differentiation, positive feedback between an ATRA-inducible signalsome complex involving many proteins including Vav1, a guanine nucleotide exchange factor, and the activation of the mitogen activated protein kinase (MAPK) cascade. The model, which was developed by integrating logical rules with kinetic modeling, was significantly smaller than previous models. However, despite its simplicity, it captured key features of ATRA induced differentiation of HL-60 cells. We identified an ensemble of effective model parameters using measurements taken from ATRA-induced HL-60 cells. Using these parameters, model analysis predicted that MAPK activation was bistable as a function of ATRA exposure. Conformational experiments supported ATRA-induced bistability. These findings, combined with other literature evidence, suggest that positive feedback is central to a diversity of cell fate programs.

*Index Terms*—Mathematical modeling, systems biology

## I. INTRODUCTION

Understanding differentiation programs is an important therapeutic challenge. Toward this challenge, lessons learned in model systems, such as the lineage-uncommitted human myeloblastic cell line HL-60, informs our analysis of more complex therapeutically important programs. Patient derived HL-60 leukemia cells have been a durable experimental model since the 1970s [2]. HL-60 undergoes cell cycle arrest and either myeloid or monocytic differentiation following stimulation; All-Trans Retinoic Acid (ATRA) induces G1/G0-arrest and myeloid differentiation in HL-60 cells, while 1,25-dihydroxy vitamin D3 (D3) induces arrest and monocytic differentiation. Commitment to cell cycle arrest and differentiation requires approximately 48 hr of treatment, during which HL-60 cells undergo two division cycles.

Sustained mitogen-activated protein kinase (MAPK) activation is a defining feature of ATRA-induced HL-60 differentiation. ATRA drives sustained MEK-dependent activation of the RAF/MEK/ERK pathway, leading to arrest and differentiation [24]. MEK inhibition results in the loss of ERK and RAF phosphorylation, and the failure to arrest and differentiate [9]. ATRA (and its metabolites) are ligands for the hormone activated nuclear transcription factors retinoic acid receptor (RAR) and retinoid X receptor (RXR) [12]. RAR/RXR activation is necessary for ATRA-induced RAF phosphorylation [9], and the formation of the ATRA-inducible signalsome complex which drives differentiation. The signalsome is composed of Src family kinases Fgr and Lyn, PI3K, c-Cbl, Slp76, and KSR, as well as IRF-1 transcription factors [5], [13]–[15], [25]. This signaling is driven by ATRA-induced expression of CD38 and the putative heterotrimeric Gq protein-coupled receptor BLR1 [4], [18]. BLR1, identified as an early ATRA (or D3)-inducible gene [23], is necessary for MAPK activation and differentiation [18]. Members of the BLR1 transcriptional activator complex, e.g. NFATc3 and CREB, are phosphorylated by ERK, JNK or p38 MAPK family members suggesting positive feedback between the signalsome and MAPK activation [22]. BLR1 overexpression enhanced RAF phosphorylation and accelerated terminal differentiation, while RAF inhibition reduced BLR1 expression and differentiation [19]. BLR1 knockout cells failed to activate RAF or differentiate in the presence of ATRA [19].

In this study, we developed a mathematical model of the key architectural feature of ATRA induced differentiation of HL-60 cells, namely positive feedback between an ATRA-inducible signalsome complex and MAPK activation. Previously Tasseff et al. hypothesized that signalsome-MAPK positive feedback was essential for ATRA-induced cell cycle arrest and differentiation [17]. We explored this hypothesis by constructing a minimal model containing only signalsome and MAPK components (Fig. 1A). The effective model was developed using a novel framework which integrated logical rules with kinetic modeling. This formulation significantly reduced the size of the model compared to the previous study of Tasseff et al., while maintaining similar model performance [17]. The effective model, despite its simplicity, captured key features of ATRA induced differentiation of HL-60 cells. We identified an ensemble of effective model parameters using measurements taken from ATRA-induced HL-60 cells. Using these parameters, model analysis predicted the bistability of MAPK activation as a function of ATRA exposure. Conformational experiments supported ATRA-induced bistability. These findings, combined with other literature evidence, suggests that positive feedback architectures are central to many cell fate programs.

**Fig. 1.**
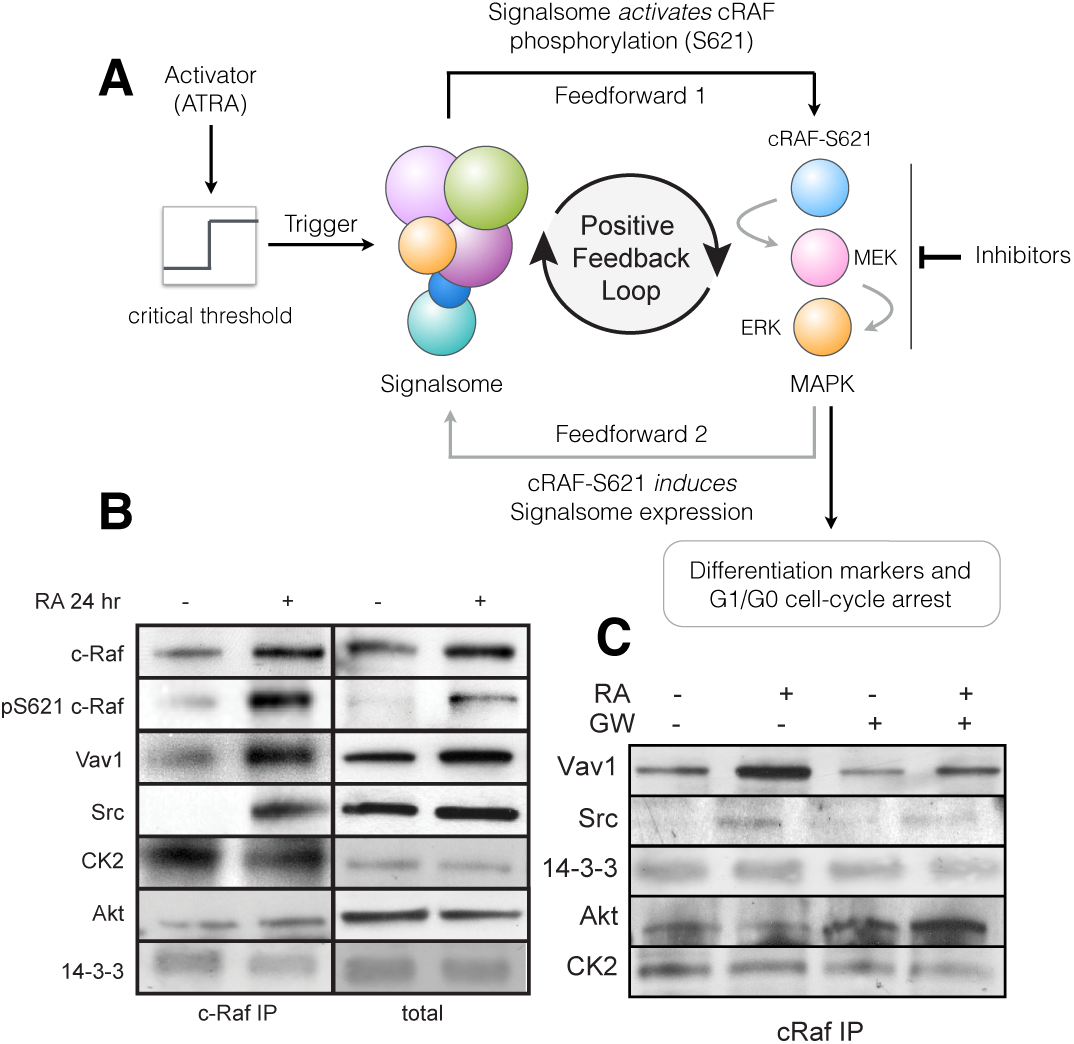
Schematic of the effective differentiation circuit. A: Above a critical threshold, ATRA activates an upstream Trigger, which induces signalsome complex formation. Signalsome activates the mitogen-activated protein kinase (MAPK) cascade which in turn drives differentiation program and signalsome formation. B: Signalsome components that interact with RAF: first column shows Western blot analysis on total RAF immunoprecipitation with and without 24 hr ATRA treatment, and the second on total lysate. C: Effect of the RAF inhibitor GW5074 on RAF interactions as determined by Western blot analysis of total RAF immunoprecipitation.

## II. RESULTS

Vav1 is a member of an ATRA-inducible signalsome complex which propels sustained MAPK activation, arrest and differentiation (Fig. 1B). We conducted immunoprecipitation studies and identified a limited number of ATRA-dependent and -independent RAF interaction partners. Of the 19 proteins sampled, Vav1, Src, CK2, Akt, and 14-3-3 precipitated with cRAF, suggesting a direct physical interaction was possible. However, only the associations between cRAF and Vav1 and Src were ATRA-inducible (Fig. 1B). Others proteins e.g., CK2, Akt and 14-3-3, generally bound cRAF regardless of phosphorylation status or ATRA treatment. Treatment with the RAF kinase inhibitor GW5074 following ATRA exposure reduced the association of both Vav1 and Src with cRAF (Fig. 1C), although the signal intensity for Src was weak. However, GW5074 did not influence the association of CK2 or 14-3-3 with cRAF, further demonstrating their independence from cRAF phosphorylation. Taken together, the immunoprecipitation and GW5074 results implicated Vav1 association to be correlated with cRAF activation following ATRA-treatment.

The model recapitulated sustained signalsome/MAPK activation following exposure to 1µM ATRA (Fig. 2A-B). An ensemble of effective model parameters was estimated by minimizing the difference between simulations and time-series measurements of BLR1 mRNA and cRAF-pS621 following the addition of 1µM ATRA using particle swarm optimization (PSO). We focused on the S621 phosphorylation site of cRAF since enhanced phosphorylation at this site is a defining characteristic of sustained MAPK activation in HL-60. Each particle in the swarm contributed a member to the parameter ensemble. The effective model captured both ATRA-induced BLR1 expression (Fig. 2A) and sustained phosphorylation of cRAF-pS621 (Fig. 2B) in a growing population of HL-60 cells. However, the effective model failed to capture the decline of BLR1 expression after 48 hr of ATRA exposure.

**Fig. 2.**
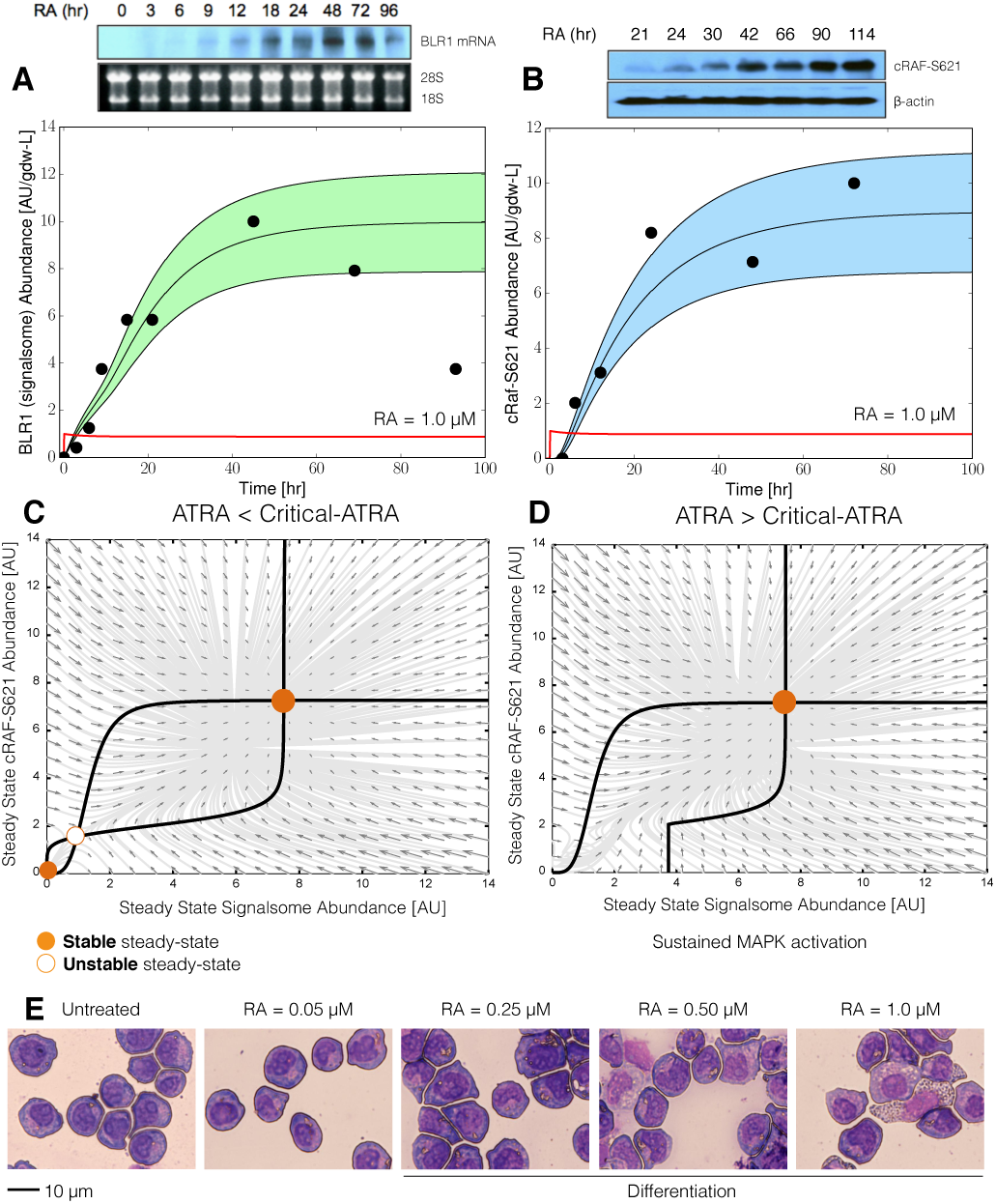
Model analysis for ATRA-induced HL-60 differentiation. A: BLR1 mRNA versus time following exposure to 1*µ*M ATRA at t = 0 hr. B: cRAF-pS621 versus time following exposure to 1*µ*M ATRA at t = 0 hr. Points denote experimental measurements, solid lines denote the mean model performance. Shaded regions denote the 99% confidence interval calculated over the parameter ensemble. C: Signalsome and cRAF-pS621 nullclines for ATRA below the critical threshold. The model had two stable steady states and a single unstable state in this regime. D: Signalsome and cRAF-pS621 nullclines for ATRA above the critical threshold. In this regime the model had only a single stable steady state. E: Morphology of HL-60 as a function of ATRA concentration (t = 72 hr).

The model was bistable with respect to ATRA induction (Fig. 2C-D). Nullcline analysis predicted two stable steady-states and a single unstable state when ATRA was present below a critical threshold (Fig. 2C). In the lower stable state, neither the signalsome nor cRAF-pS621 were present (thus, the differentiation program was deactivated). However, at the high stable state, both the signalsome and cRAF-pS621 were present, allowing for sustained activation and differentiation. Interestingly, when ATRA was above a critical threshold, only the activated state was accessible (Fig. 2D). To test these findings, we first identified the ATRA threshold. We exposed HL-60 cells to different ATRA concentrations for 72 hr (Fig. 2E). Morphological changes associated with differentiation were visible for ATRA ≥ 0.25 *µ*M, suggesting the critical ATRA threshold was near this concentration.

Next, we tested that a cell locked into an activated state remained activated following ATRA withdraw. Sustained activation resulted from reinforcing feedback between the signalsome and the MAPK pathway. After activation, if we inhibited or removed elements from the effective circuit we expected the siganlsome and MAPK signals to decay. We simulated ATRA induced activation in the presence of kinase inhibitors, and without key circuit elements (Fig. 3). Consistent with experimental results using multiple MAPK inhibitors, ATRA activation in the presence of MAPK inhibitors lowered the steady-state value of signalsome (Fig. 3A). In the presence of BLR1, the signalsome and cRAF-pS621 signals were maintained following ATRA withdraw (Fig. 3B, blue). On the other hand, BLR1 deletion removed the ability of the circuit to maintain a sustained MAPK response following the withdraw of ATRA (Fig. 3B, gray). Lastly, washout experiments in which cells were exposed to 1*µ*M ATRA for 24 hr, and then transferred to fresh media without ATRA, confirmed the persistence of the self sustaining activated state for up to 144 hr (Fig. 3C).

**Fig. 3.**
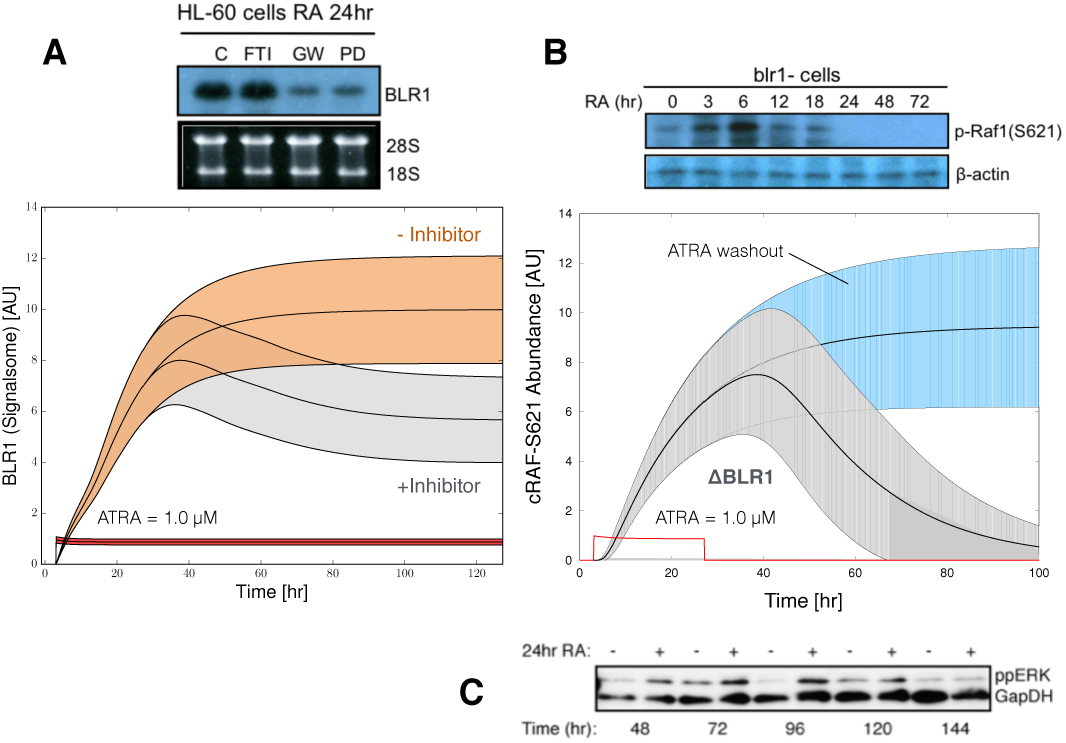
Model simulation follow exposure to 1*µ*M ATRA. A: BLR1 mRNA versus time with and without MAPK inhibitor. B: cRAF-pS621 versus time following pulsed exposure to 1*µ*M ATRA with and without BLR1. Solid lines denote the mean model performance, while shaded regions denote the 99% confidence interval calculated over the parameter ensemble. C: Western blot analysis of phosphorylated ERK1/2 in ATRA washout experiments. Experimental data in panels A and B were reproduced from Wang and Yen [19], data in panel C is reported in this study.

## III. DISCUSSION

We presented an effective model of ATRA-inducible differentiation of HL-60 cells which encoded positive feedback between the ATRA-inducible signalsome complex and the MAPK pathway. Despite its simplicity, the model captured key features of the ATRA induced differentiation such as sustained MAPK activation, and bistability with respect to ATRA exposure. We also reported a new ATRA-inducible component of the signalsome, Vav1. Vav1 is a guanine nucleotide exchange factor for Rho family GTPases that activate pathways leading to actin cytoskeletal rearrangements and transcriptional alterations [10]. The Vav1/RAF association correlated with RAF activity, was ATRA-inducible and decreased after treatment with GW5074. The presence of Vav1 in RAF/Grb2 complexes has been shown to correlate with increased Raf activity in mast cells [16]. Furthermore, studies on Vav1 knockout mice demonstrated that the loss of Vav1 resulted in deficiencies of ERK signaling for both T-cells as well as neutrophils [6], [8]. While its function in the signalsome is unclear, Vav1 has been shown to associate with a Cbl-Slp76-CD38 complex in an ATRA-dependent manner; furthermore, transfection of HL-60 cells with Cbl mutants that fail to bind CD38, yet still bind Slp76 and Vav1, prevented ATRA-induced MAPK activation [15]. Thus, interaction of Cbl-Slp76-Vav1 and CD38 appears to be required for transmission of the ATRA signal by the signalsome.

Several engineered, or naturally occurring systems involved in cell fate decisions incorporate positive feedback and bistability [7]. One of the most well studied cell fate circuits is the Mos mitogen-activated protein kinase cascade in *Xenopus* oocytes. This cascade is activated when oocytes are induced by the steroid hormone progesterone [21]. The MEK-dependent activation of p42 MAPK stimulates the accumulation of the Mos oncoprotein, which in turn activates MEK, thereby closing the feedback loop. This is similar to the differentiation circuit presented here; ATRA drives signalsome which activates MAPK, cell-cycle arrest, differentiation and signalsome. Thus, while HL-60 and *Xenopus* oocytes are vastly different biological models, they share similar cell fate decision architectures. Other unrelated cell fate decisions such as programmed cell death have also been suggested to be bistable [1]. Still more biochemical networks important to human health, for example the human coagulation or complement cascades, also feature strong positive feedback elements [11]. Thus, while positive feedback is sometimes not desirable in man made systems, it may be at the core of a diverse variety of cell fate programs and other networks important to human health.

Model performance was impressive given its limited size. However, there were several issues to explore further. First, there was likely missing connectivity in the effective differentiation circuit. Decreasing BLR1 expression with simultaneously sustained cRAF-pS261 activation was not captured by the current network architecture. This suggested that signalsome, once activated, had a long lifetime as decreased BLR1 expression did not impact cRAF-pS261 abundance. We could model this by separating siganlsome formation into an inactive precursor pool that is transformed to a long-lived activated siganlsome by MAPK activation. We should also explore adding additional downstream biological modules to this skeleton model, for example the upregulation of reactive oxygen markers such as p47Phox or cell cycle arrest components to capture the switch from an actively proliferating population to a population in G0-arrest. Next, the choice of max/min integration rules or the particular form of the transfer functions could also be explored. Integration rules other than max/min could be used, such as the mean or the product, assuming the range of the transfer functions is always *f* ∈ [0, 1]. Alternative integration rules might have different properties which could influence model identification or performance. For example, a mean integration rule would be differentiable, allowing derivative-based optimization approaches to be used. The form of the transfer function could also be explored. We choose hill-like functions because of their prominence in the systems and synthetic biology community. However, many other transfer functions are possible.

## IV. MATERIALS AND METHODS

*Effective model equations:* The model was constructed using the hybrid approach of Wayman et al. [20]. The abundance of species *i* (*x_i_*) is governed by:

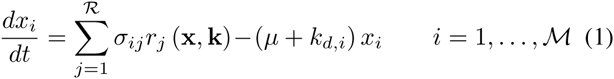

where 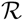 and 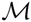 denotes the number of reactions and species in the model, *µ* denotes the specific growth rate and *k_d,i_* denotes a degradation constant for species *i*. The *µx_i_* term, which accounts for dilution due to cell growth, decays to zero following ATRA exposure. The quantity *σ_ij_* denotes the stoichiometric coefficient for species *i* in reaction *j*, *r_j_* (x*, ε,* k) denotes the rate of reaction *j*, and k 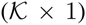 denotes the unknown kinetic parameter vector. If *σ_ij_ >* 0, species *i* is produced by reaction *j*, if *σ_ij_* < 0, species *i* is consumed by reaction *j*, while *σ_ij_* =0 indicates species *i* is not connected with reaction *j*. Species balances were subject to the initial conditions x (*t_o_*)= x*_o_*. The reaction rate was written as the product of a kinetic term 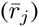 and a control term (*v_j_*), 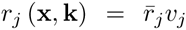. In this study, we used either zero-or first-order kinetics. The control term 0 ≤ *v_j_* ≤ 1 depended upon the combination of factors which influenced rate process *j*. For each rate, we used a rule-based approach to select from competing control factors. If rate j was influenced by 1,&,*m* factors, we modeled this relationship as 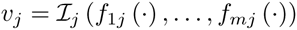 where 0 ≤ *f_ij_* (·) ≤ 1 denotes a regulatory transfer function quantifying the influence of factor *i* on rate *j*. The function 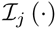 is an integration rule which maps the output of regulatory transfer functions into a control variable. Each regulatory transfer function took the form:

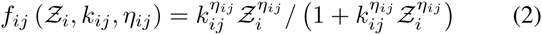

where 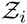 denotes the abundance factor *i*, *k_ij_* denotes a gain parameter, and *η_ij_* denotes a cooperativity parameter. In this study, we used 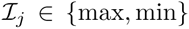 [20]. If a process has no modifying factors, *v_j_* =1.

*Estimation of model parameters:* Model parameters were estimated by minimizing the squared difference between simulations and experimental data set *j*:

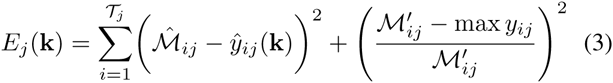

The terms 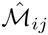 and 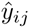 denote scaled experimental observations and simulation outputs at time *i* from training set *j*, where 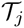 denoted the number of time points for data set *j*. The first term in Eqn. (3) quantified the relative simulation error. We used immunoblot intensity measurements for model training. Thus, we trained the model on the *relative* change between bands within each data set. Suppose we have the intensity of species *x* at time {*t*_1_, *t*_2_, …, *t_n_*} in condition *j*. The scaled value 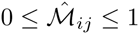 is given by:

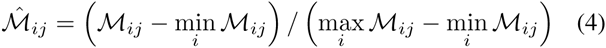

where 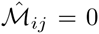 and 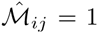 describe the lowest (highest) intensity bands. A similar scaling was used for the simulation output. The second term in the objective function ensured a realistic concentration scale was estimated by the model. We set the highest intensity band to 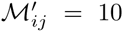 [AU] for all simulations. We minimized the total model residual ∑*_j_ E_j_* using particle swarm optimization (PSO). All code was implemented in the Octave programming language, and is available for download from http://www.varnerlab.org.

*Cell culture and treatment:* Human myeloblastic leukemia cells (HL-60 cells) were grown in a humidified atmosphere of 5% CO_2_ at 37^o^C and maintained in RPMI 1640 from Gibco (Carlsbad, CA) supplemented with 5% heat inactivated fetal bovine serum from Hyclone (Logan, UT) and 1× antibiotic/antimicotic (Gibco, Carlsbad, CA). Cells were cultured in constant exponential growth [3]. Experimental cultures were initiated at 0.1 × 10^6^ cells/mL 24 hr prior to ATRA treatment; if indicated, cells were also treated with GW5074 (2*µ*M) 18 hr before ATRA treatment. For the cell culture washout experiments, cells were treated with ATRA for 24 hr, washed 3x with prewarmed serum supplemented culture medium to remove ATRA, and reseeded in ATRA-free media as described. Western blot analysis was performed at incremental time points after removal of ATRA.

*Chemicals:* All-Trans Retinoic Acid (ATRA) from Sigma-Aldrich (St. Louis, MO) was dissolved in 100% ethanol with a stock concentration of 5mM, and used at a final concentration of 1*µ*M (unless otherwise noted). The cRAF inhibitor GW5074 from Sigma-Aldrich (St. Louis, MO) was dissolved in DMSO with a stock concentration of 10mM, and used at a final concentration of 2*µ*M. HL-60 cells were treated with 2*µ*M GW5074 with or without ATRA (1*µ*M) at 0 hr. This GW5074 dosage had a negligible effect on the cell cycle distribution, compared to ATRA treatment alone.

*Immunoprecipitation and western blotting:* Approximately 1.2 × 10^7^ cells were lysed using 400*µ*L of M-Per lysis buffer from Thermo Scientific (Waltham, MA). Lysates were cleared by centrifugation at 16,950 × g in a micro-centrifuge for 20 min at 4^o^C. Lysates were pre-cleared using 100*µ*L protein A/G Plus agarose beads from Santa Cruz Biotechnology (Santa Cruz, CA) by inverting overnight at 4^o^C. Beads were cleared by centrifugation and total protein concentration was determined by a BCA assay (Thermo Scientific, Waltham, MA). Immunoprecipitations were setup by bringing lysate to a concentration of 1g/L in a total volume of 300*µ*L (M-Per buffer was used for dilution). The anti-RAF antibody was added at 3*µ*L. A negative control with no bait protein was also used to exclude the direct interaction of proteins with the A/G beads. After 1 hr of inversion at 4^o^C, 20*µ*L of agarose beads was added and samples were left to invert overnight at 4^o^C. Samples were then washed three times with M-Per buffer by centrifugation. Finally proteins were eluted from agarose beads using a laemmli loading buffer. Eluted proteins were resolved by SDS-PAGE and Western blotting. Total lysate samples were normalized by total protein concentration (20*µ*g per sample) and resolved by SDS-PAGE and Western blotting. Secondary HRP bound antibody was used for visualization. All antibodies were purchased from Cell Signaling (Boston, MA) with the exception of *α*-p621 RAF which was purchased from Biosource/Invitrogen (Carlsbad, CA), and *α*-CK2 from BD Biosciences (San Jose, CA).

*Morphology assessment:* Untreated and ATRA-treated HL-60 cells were collected after 72 hr and cytocentrifuged for 3 min at 700 rpm onto glass slides. Slides were air-dried and stained with Wrights stain. Slide images were captured at 40X (Leica DM LB 100T microscope, Leica Microsystems).

